# Triflic acid treatment enables LC-MS/MS analysis of insoluble bacterial biomass

**DOI:** 10.1101/367599

**Authors:** Ana Y. Wang, Peter S. Thuy-Boun, Gregory S. Stupp, Andrew I. Su, Dennis W. Wolan

**Affiliations:** Department of Molecular Medicine and Department of Integrative Structural and Computational Biology, The Scripps Research Institute, 10550 North Torrey Pines Road, La Jolla, CA 92037, USA

**Keywords:** triflic acid, membrane proteins, microbiome, metaproteomics, MudPIT, ComPIL

## Abstract

The lysis and extraction of soluble bacterial proteins from cells is a common practice for proteomics analyses, but insoluble bacterial biomasses are often left behind. Here, we show that with triflic acid treatment, the insoluble bacterial biomass of Gram^-^ and Gram^+^ bacteria can be rendered soluble. We use LC-MS/MS shotgun proteomics to show that bacterial proteins in the soluble and insoluble post-lysis fractions differ significantly. Additionally, in the case of Gram^-^ *Pseudomonas aeruginosa*, triflic acid treatment enables the enrichment of cell envelope-associated proteins. Finally, we apply triflic acid to a human microbiome sample to show that this treatment is robust and enables the identification of a new, complementary subset of proteins from a complex microbial mixture.

## INTRODUCTION

Analysis of intracellular proteins by shotgun proteomics requires cells to be lysed and the contents proteolyzed for LC-MS/MS.^1–4^ Simple cell disruption methods (e.g., detergents, sonication) are sufficient for the full lysis and solubilization of human cells. However, for single bacterial species and complex microbial mixtures (e.g., microbiomes), simple methods of cell disruption yield soluble protein and leave behind large, insoluble, proteolytically resistant pellets. More effective methods of microbial cell disruption^5–7^ utilizing chaotropes,^8,9^ detergents,^10^ bead beating,^11^ sonication,^12–15^ freeze-thawing,^16,17^ aqueous acid/base treatments,^18,19^ organic extraction,^20^ or a combination thereof^3,4,21–38^ have been applied to recalcitrant microbiota isolated from terrestrial, aquatic, and gastrointestinal systems in the past. Our attempts to adapt many of these methods to microbiome samples yielded familiar results, including soluble protein solutions and large insoluble, proteolytically resistant pellets. The size of these remaining pellets lead us to question whether residual protein is trapped in this insoluble matrix and overlooked by downstream analyses. We reasoned that the pellets’ resistance to solubilization and proteolysis could partly be due to carbohydrate-rich bacterial macromolecules that include peptidoglycans, lipopolysaccharides, lipotechoic acids, and other cell-envelope constituents that are absent in human cells.^39^ We therefore sought a method to fragment and separate carbohydrate macromolecules from insoluble bacterial pellets without destroying any putatively associated pellet proteins.

Trifluoromethanesulfonic acid (triflic acid, TA) is used to fully remove glycans from *N*-linked, *O*-linked, and glycosaminoglycan-containing glycoproteins while preserving peptide integrity. TA can also de-glycosylate *N-*linked glycoproteins, leaving behind a single *N*-linked monosaccharide under highly controlled conditions.^40,41^ This reactivity was recently utilized by the Wu group to map protein *N*-glycosylation sites of the soluble yeast proteome by LC-MS/MS.^42^ We hypothesized that if insoluble microbial pellets are composed largely of carbohydrates, TA could remove these molecules, enabling proteomic analysis of any putatively associated proteins.

Three species with distinct cell envelope morphologies were selected for TA treatment or sonication and downstream LC-MS/MS analysis: (1) Gram^-^ *Pseudomonas aeruginosa*, with a peptidoglycan layer between two membranes; (2) Gram^+^ *Bacillus subtilis*, with an exposed peptidoglycan layer; and (3) human Jurkat cells - a peptidoglycan-less control. We then TA-treated a human distal gut microbiome sample for LC-MS/MS analysis to evaluate the robustness of this treatment for complex microbial mixtures.

## MATERIALS AND METHODS

### Sonication-based proteome preparation of single species

Overnight cultures of *Pseudomonas aeruginosa* or *Bacillus subtilis* (grown in 5 mL 2XYT media at 37 °C) and human Jurkat cells (grown in RPMI-1640 Medium with 10% heat-inactivated fetal bovine serum) were divided into 100-mg wet pellets. The cells were suspended in 500 μL PBS, pH 7.4 and sonicated in a Qsonica Q700 sonicator with cup horn attachment at 25% amplitude for 10 min at 4 °C. Insoluble cellular debris was removed by centrifugation at 10,000 x*g* for 5 min at room temperature, and the supernatant was carefully removed and filtered through a PD MiniTrap G-25 size exclusion column (GE Life Sciences, #28918007) to remove residual, low molecular-weight contaminants. The soluble protein content in the flow-through (samples designated as A-supernatant, see Supporting Information, Figure 1 for details) was then quantified using the Pierce^TM^ BCA Protein Assay Kit (Thermo Fisher, #23225).

**Figure 1.**
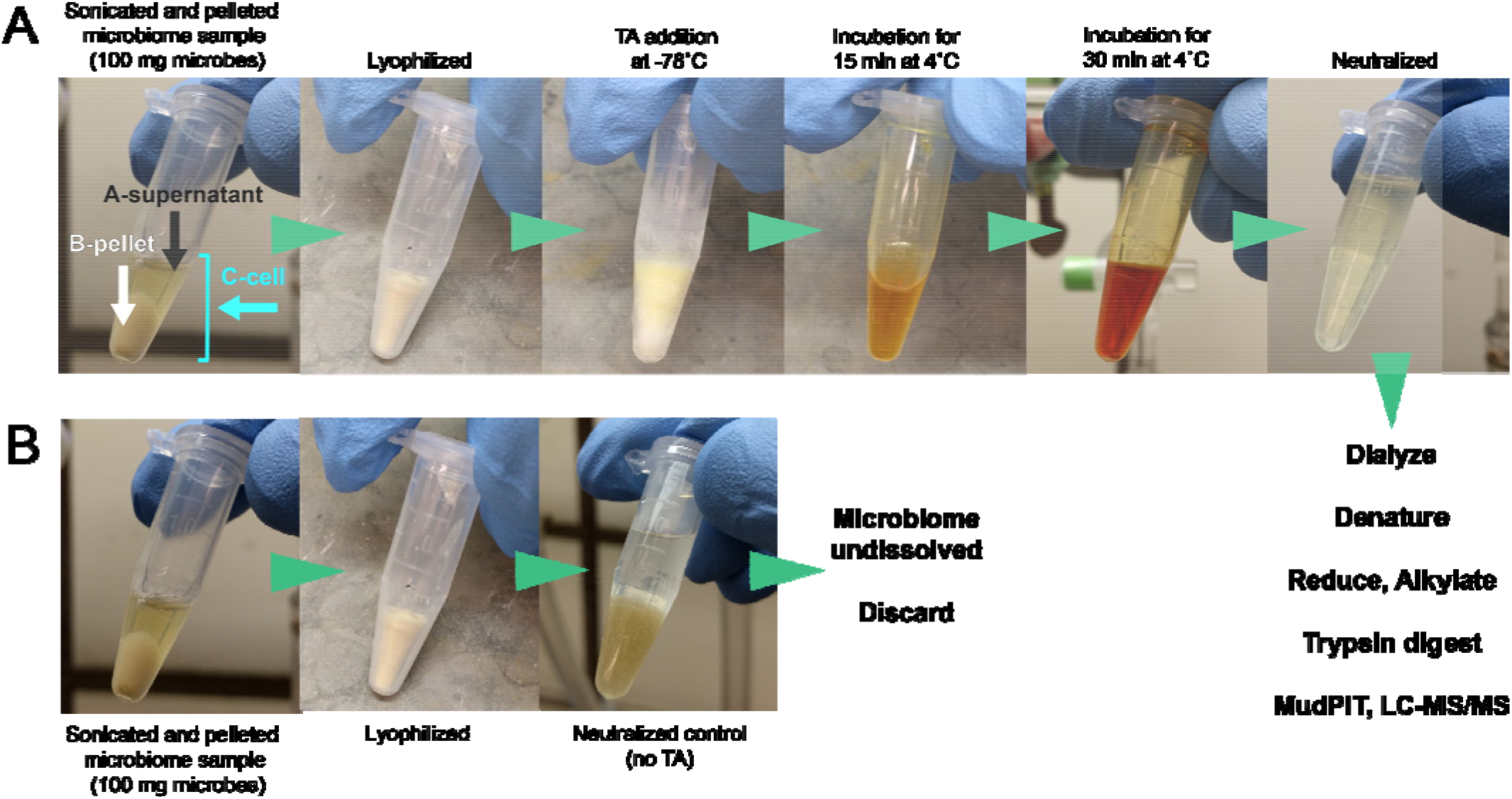
Simplified proteome preparation and analysis schematic. (A) Triflic acid (TA) treatment of a microbio sample. Note that whole bacterial cells and pellets dissolve to give a red-orange solution with TA treatment. T mixture remains soluble after neutralization (far right frame). (B) Control: treatment of lyophilized non-TA tre microbiome sample with neutralization solution. Note the remaining undissolved microbial biomass after treatment with neutralizing solution alone.

### Triflic acid proteome preparation of single species

To prepare the TA-treated samples (samples designated as C-cell), 100-mg (wet pellet) aliquots of *P. aeruginosa, B. subtilis*, and Jurkat cells were suspended in 500 μL PBS, pH 7.4, frozen, and lyophilized overnight. 200 μL of 1:9 toluene:TA (TCI, #T0751, CAS[1493–13-6]) was added to each lyophilized whole-cell sample at –80 °C (dry ice/acetone bath). Tubes containing the solutions were brought to 4 °C and gently agitated for 30–60 min with intermittent venting until the pellet was fully dissolved. Upon complete solubilization, the solution was refrozen at –80 °C (dry ice/acetone bath). The acid was quenched with 600 μL of a cold solution of 1:1:3 water:methanol:pyridine (neutralizing solution), and the tubes were inverted several times with frequent venting. The neutralized solution was diluted to 10 mL with 50% methanol/water and concentrated to 1 mL with a 3K MWCO centrifugal concentrator (Fisher Scientific, Amicon Ultra-15 3K, UFC900324). The sample was diluted to 10 mL with PBS, pH 7.4 and re-concentrated with a 3K MWCO concentrator. The remaining solution and protein were transferred to a microcentrifuge tube and lyophilized. The protein solid was resuspended in 200 μL of 5 M urea in 100 mM Tris, pH 8.0 and sonicated until the protein dissolved (see Figure S1 for details). The protein concentration was quantified using the BCA assay.

The remaining insoluble, post-sonication pellets from the *B. subtilis* and *P. aeruginosa* proteome preparations were resuspended in PBS pH 7.4, frozen, lyophilized, and TA-treated with the same methods as described for the C-cell preparations (samples are designated as B-pellet, Figure S1).

### Sonication and triflic acid preparations of microbiome samples

Fecal samples were suspended in PBS and mixed by vortexing until full suspension and uniformity. Samples were centrifuged at 100 x*g* for 1 min, and the cloudy supernatant was filtered through a 70-µm nylon cell strainer (Fisher Scientific, Falcon^TM^ #08–771-2) to remove large particles. The microbial cells were pelleted by centrifugation at 8,000 x*g* for 5 min, rinsed twice with PBS, pH 7.4, and divided into 100-mg cell pellets (wet weight) for each sample. For C-cell preparation, microbial cell pellets were suspended in PBS, pH 7.4, frozen, and lyophilized. 300 µL of 1:9 toluene:TA was added to each sample at –80 °C (dry ice/acetone bath). The sample was raised to 4 °C and gently agitated for 30–60 min at 4 °C with intermittent venting. After the cell pellet was completely liquefied, the solution was frozen at –80 °C (dry ice/acetone bath) and quenched with 900 µL of a cold solution of 1:1:3 water:methanol:pyridine. The tubes were inverted with frequent venting until the frozen pellet was dissolved. This solution was diluted to 10 mL with 50% methanol/water, concentrated to 1 mL with a 3K MWCO centrifugal concentrator, and then diluted and re-concentrated twice more with 100 mM Tris HCl, pH 8.0. The remaining protein suspension was transferred to a 1.5 mL microcentrifuge tube, frozen, and lyophilized. The protein was resuspended in 5 M urea in 100 mM Tris HCl, pH 8.0 and sonicated until completely dissolved, and the protein concentration was quantified using the BCA assay (Figure S1). For B-pellet preparation, the pellet remaining after sonication was TA-treated, and protein was extracted similarly to single-species B-pellet samples (Figure S1). For the microbiome A-supernatant sample preparation, soluble proteins were collected after microbiome sonication, centrifugation, and size exclusion purification (PD MiniTrap G-25) similarly to single-species A-supernatant samples (Figure S1). However, an additional step was required that included the collected supernatant to be frozen, lyophilized overnight, and then TA-treated (as for B-pellet and C-cell microbiome fractions).

### Preparation of samples for LC-MS/MS analysis

100 μg protein per sample suspended in 120 μL 100 mM Bicine, 5 M urea, pH 8.0 were treated with tris(2-carboxyethyl)phosphine (TCEP) (Thermo Fisher, Pierce, 20490) to a final concentration of 5 mM and incubated for 15 min, followed by chloroacetamide (Sigma-Aldrich, 201-174-2, CAS[79-07-2) treatment at a final concentration of 25 mM for 15 min in the dark. The mixture was then diluted to a final volume of 500 μL with trypsin buffer (100 mM Tris HCl, pH 8.0 with 1mM calcium chloride), and 2 μg of trypsin (Promega, V5111) was incubated with each sample overnight at 37 °C under agitation. After incubation, samples were treated with 25 μL formic acid and the samples were centrifuged at 12,000 x*g* for 5 min. 475 μL of the topmost liquid layer were separated and stored at –20 °C.

### LC-MS/MS MudPIT data collection

The trypsin-digested peptides of single-species samples were loaded onto a biphasic column (250 μm fused silica, Thomas Scientific 2713X55), with 3 cm of 5 μm Aqua C18 resin (Phenomenex, 00G-4299-E0) followed by 3 cm of Partisphere strong cation exchange (SCX) resin. The analytical column was made by pulling 100 μm fused silica (Thomas Scientific, 2713X83) to a 5 μm tip with a micropipette puller (Sutter Instrument Company, Model P-2000) and packed with 10 cm of 5 μm Aqua C18 resin. The sample and analytical columns were joined using a zero-dead volume union (Waters).

Multidimensional protein identification technology (MudPIT)^43,44^ tandem mass spectrometry (LC-MS/MS) was performed using a Thermo Finnigan LTQ attached to an Agilent 1200 series quaternary pump. The pump was set at 200 μL/min, and peptides were eluted at a rate of 250 nL/min, using a 6-step MudPIT program. The first step began with 2.5 min of 100% buffer A (95% H_2_O, 5% acetonitrile, 0.1% formic acid), followed by 30 min of 15% buffer B (20% H2O, 80% acetonitrile, 0.1% formic acid) and 85% buffer A. Buffer B was then increased over 20 min to 100% before ending with 20 min of 100% buffer A. Each salt step began with 1 min of 100% buffer A, followed by a 4-min salt pulse with various concentrations of buffer C (500 mM ammonium acetate, 95% H_2_O, 5% acetonitrile, 0.1% formic acid), then 5 min of 100% buffer A, followed by a 105-min gradient from 5%-65% buffer B, and ending with 5 min of 100% buffer A. The salt pulses of buffer C were as follows: 10%, 25%, 50%, 80%, 100%, with buffer A as the diluent. The MS was operated with the following settings: MS1 mass range of 300-2,000, 5 precursor ions selected for tandem MS per scan cycle, intensity threshold of 500 for triggering MS2, relative collision energy 35%, and dynamic exclusion set to exclude after two times if occurs within 30 sec with an exclusion duration of 20 sec.

For microbiome samples, 40 µL of the peptide solution from “Proteome sample preparation for LC-MS/MS” was dried using a SpeedVac concentrator and desalted using 10 µL ZipTips C 18 (Millipore). The resulting peptide solution dried using the SpeedVac concentrator and was analyzed with a Thermo Fisher Orbitrap Fusion^TM^ Tribrid^TM^ mass spectrometer coupled to a nLC 1000 system in the Proteomics Core at TSRI Florida.

The dry peptides were reconstituted in 40 µL of 0.1% formic acid in water, and 10 µL of the peptide solution were used for LC-MS/MS analysis. Peptides were on-line eluted onto an analytical reverse phase column (0.075 × 150 mm Acclaim PepMap RLSC nano Viper, Thermo Fisher) at 300 nL/min with the following gradients, using buffer A (0.1% formic acid in H_2_O) as the diluent: 5–25% buffer B (80% acetonitrile, 0.1% formic acid, 20% H_2_O) in 160 min, 25–44% buffer B in 80 min, 44–80% buffer B in 10 sec, 80% buffer B for 5 min, 80–5% buffer B in 10 sec, and 5% B for 20 min. The MS was operated with the following settings: MS1 scan range of 380–1400 m/z with a mass tolerance of 10 ppm and 120K resolution using Orbitrap detection, data-dependent MS/MS mode at the maximum speed with precursor priority of the most intense ions, HCD fragmentation with normalized collision energy of 30%, 1.0E4 AGC Target intensity threshold for MS2, 2–8 charge states included in screening parameters, a mass resolution of 30K for MS2, and dynamic exclusion set to exclude after two times if occurs within 30 sec with an exclusion duration of 20 sec. All LC-MS/MS data have been deposited to the ProteomeXchange Consortium (http://proteomecentral.proteomexchange.org) via the PRIDE partner repository with the project accession identifier PXD009004.^45^

### Peptide and protein identification

For the MS data of single-species samples, rawXtract 1.9.9.2 was used to extract precursor and fragmentation ion data in MS2 format from Xcalibur RAW files. Integrated Proteomics Pipeline (IP2, Integrated Proteomics Applications, Inc. San Diego, CA) was utilized for protein identification and quantification analyses, using PROLUCID/Sequest^46,47^ and DTASelect.^47,48^ Tandem mass spectra were extracted into the MS2 file format from the Xcalibur RAW files via RawXtract 1.9.9.2 (http://fields.scripps.edu/yates/wp/?page_id=17). The MS2 spectra were scored against the protein library of each respective species from Uniprot proteome IDs, UP000001570 (*Bacillus subtilis* 168), UP000002438 (*Pseudomonas aeruginosa* PA01), and UP000005640 (*Homo sapiens*), which include the reversed sequence for each entry in the original database,^49^ using PROLUCID/Sequest. The settings for peptide scoring include: (1) differential modifications of oxidized methionine (+15.9949 Da) and GlcNAc-tagged asparagine (+203.0794 Da), (2) a static modification for alkylated cysteine residues (+57.02146 Da), (3) precursor/peptide mass tolerance of 3 Da (4) and a fragment mass tolerance of 600 ppm. The search space included half- and fully tryptic peptide candidates with two missed cleavage events and required a minimum of two peptides per protein. The ProLUCID search results were assembled and filtered using the DTASelect program. Spectral counts (SC) determined from this pipeline were used as the basis of our quantitative analyses. Venn diagrams were generated based on the presence or absence of one or more SC in each protein.

The human microbiome samples’ MS2 spectra were scored using Blazmass 0.9993 against the peptides of the Comprehensive Protein Identification Library (ComPIL) database.^50^ Blazmass and ComPIL source code are open source (https://github.com/sandipchatterjee/blazmass_compil). Settings for peptide scoring include a variable modification of oxidized methionine (+15.9949 Da), a static modification for alkylated cysteine residues (+57.02146 Da), a precursor mass tolerance of 10 ppm, and a fragmentation ion tolerance of 50 ppm. DTASelect 2.1.3 (http://fields.scripps.edu/yates/wp/?page_id=17) was used for filtering with the requirements of two peptides per protein and a protein FDR of 1%.

The source code for this analysis is available online (https://github.com/stuppie/triflic/tree/master/H1). Protein clustering, cluster taxonomy, and gene ontology (GO) term annotations were performed as previously described.^50–52^ Protein loci were mapped to protein clusters with ComPIL using a sequence identity threshold of 70%. A protein cluster was annotated with all GO terms associated with any domain for all possible proteins in that cluster. Any GO terms that were parents (“is a” or “part of” relationships) of other GO terms in that protein cluster were removed. Annotations were generated from InterProScan v5 (version 5.8–49.0).

### Differential analysis of detected proteins

Protein differential analysis was performed using DESeq2 1.16.1, which allows for testing for differential expression using SC data and provides methods designed for taking into account over-dispersion, features with low counts, and experiments with low numbers of replicates.^53^ This program has previously been applied to proteomic studies.^51,52,54^ Briefly, count data is modeled using the negative binomial distribution, and the mean-variance relationship is estimated. Variance is estimated using an information sharing approach, whereby a single feature’s (or protein cluster’s) variance is estimated by taking into account variances of other protein clusters measured in the same experiment. Feature significance calling and ranking is performed using estimated effect sizes, which takes into account the logarithmic fold change (LFC) of a protein locus between enriched and unenriched samples and the noisiness of the LFC estimate. Replicates were not merged, but were treated as replicates for the differential analysis. The PCA plot was generated using DESeq2 on the variance-stabilized (*i.e*.,transformed) SC for each protein or protein locus. All volcano plots were created Prism v5 (GraphPad, Inc.). SC calculated for the clusters are the sum of the SC of all proteins within the cluster identified by LC-MS/MS. Tables S1–S16 highlight the top 10 most positively or negatively differentially detected proteins among sample preparation methods. A full list of all detected proteins is provided in the Supporting Information Dataset.

### GO Slim terms for microbiome sample analysis

Each protein locus was mapped to Gene Ontology (GO)^55,56^ terms associated with any domain of all possible proteins within each locus identified by LC-MS/MS. A GO Slim mapping file was generated by downloading GO mapping files from Uniprot-GOA (released 2017–06-06; ftp://ftp.ebi.ac.uk/pub/databases/GO/goa/UNIPROT/goa_uniprot_gcrp.gaf.gz) and the Gene Ontology (release 2017-06-19, http://purl.obolibrary.org/obo/go.obo). A simplified version of the GO (GO slim) was created using Owltools (https://github.com/owlcollab/owltools), and it consisted of the terms within the Cellular Component (CC) ontology, including membrane (GO:0016020), cell (GO:0005623), cytoplasm (GO:0005737), organelle (GO:0043226), cell periphery (GO:0071944), and the ontology CC (GO:0005575) itself. The resulting dictionary generated contains the Uniprot ID of each protein as the key, and the value is the list of GO CC terms annotated to that protein cluster.

### Assignment and quantification of cellular component GO annotations

CC annotations associated with each protein cluster were retrieved from the generated dictionary, and the total SC of peptides within each cluster were summed. The count contributed by each protein cluster to a CC annotation was calculated by dividing the total sum of the SC of each cluster by the number of CC annotations associated with that cluster, which controls for inflation of values due to annotation biases (*i.e*. more- versus less-studied/annotated proteins).

The cellular localizations in Figure 2C were derived from the CC annotations we retrieved for the protein clusters identified by LC-MS/MS. Four categories were created by binning together the CC terms as follows: 1) *Membrane & Cell Envelope*: membrane (GO:0016020), cell periphery (GO:0071944); 2) *Cytosol*: cytoplasm (GO:0005737); 3) *Cell*: cell (GO:0005623), organelle (GO:0043226), cellular component (GO:0005575); and 4) *No CC Annotation*: no CC annotations listed for the protein.

## RESULTS AND DISCUSSION

*P. aeruginosa, B. subtilis*, Jurkat cells, and a healthy human microbiome sample were partitioned into two sets. Samples in the first set were lysed by sonication^52^ (A-supernatant), clarified by centrifugation, and prepared for LCMS/MS analysis (Figure 1). While Jurkat cells fully dissolved, bacterial samples yielded yellow supernatants and insoluble pellets. The remaining *P. aeruginosa, B. subtilis*, and microbiome pellets (B-pellet) were TA-treated and prepared for LC-MS/MS analysis.^43,44^ The second set of samples were TA-treated directly and prepared for LCMS/MS (C-cell) (Figure 1). TA treatment completely dissolved whole bacterial cells and the previously insoluble bacterial pellets left after sonication. Importantly, TA treatment yielded material that is readily solubilized after trypsin digestion or detergent/chaotrope treatment, post-workup. Substantial differences in the compositions of the detected proteomes of each treatment group were observed by LC-MS/MS (see Supporting Information Dataset). Here, we highlight the proteins with the greatest differences in abundance due to proteome preparation methods.

### TA treatment of *P. aeruginosa* markedly increases cell envelope-associated protein detection

There were striking differences between A-supernatant, B-pellet, and C-cell datasets for *P. aeruginosa*. 2360 proteins were collectively identified, with 1118 shared proteins and 234–242 proteins uniquely detected by specific treatments (Figure 2). Comparison of A-supernatant and C-cell showed that the detected quantities of 78 proteins changed significantly with an absolute log_2_ fold-change (|log_2_FC|) >1 (Figure 3A). 60/78 proteins experienced enrichments of log_2_FC 1.58 to 11.8 with TA treatment. Of the 10 most-shifted proteins, 9 are cell envelope-associated (Table S1). Among the most positively enriched proteins were two envelope-associated, uncharacterized proteins (Q9HVI2, Q9HVK6), large-conductance mechanosensitive channel (Q9HVH7), and outer membrane (OM) porin F (P13794), with a log_2_FC of 11.8 and 7.86, 7.42, and 7.08, respectively. Conversely, 18 proteins were negatively enriched in the C-cell dataset, with 5 of the 10 most shifted proteins annotated as cytoplasmic, two as periplasmic, and three with unknown localizations (Supporting Information, Table 2). These negatively enriched cytoplasmic proteins include formyltetrahydrofolate deformylase (Q9HW87) and pterin-4-alpha-carbinolamine dehydratase (P43335), with log_2_FC values of –5.74 and –4, respectively.

**Figure 2.**
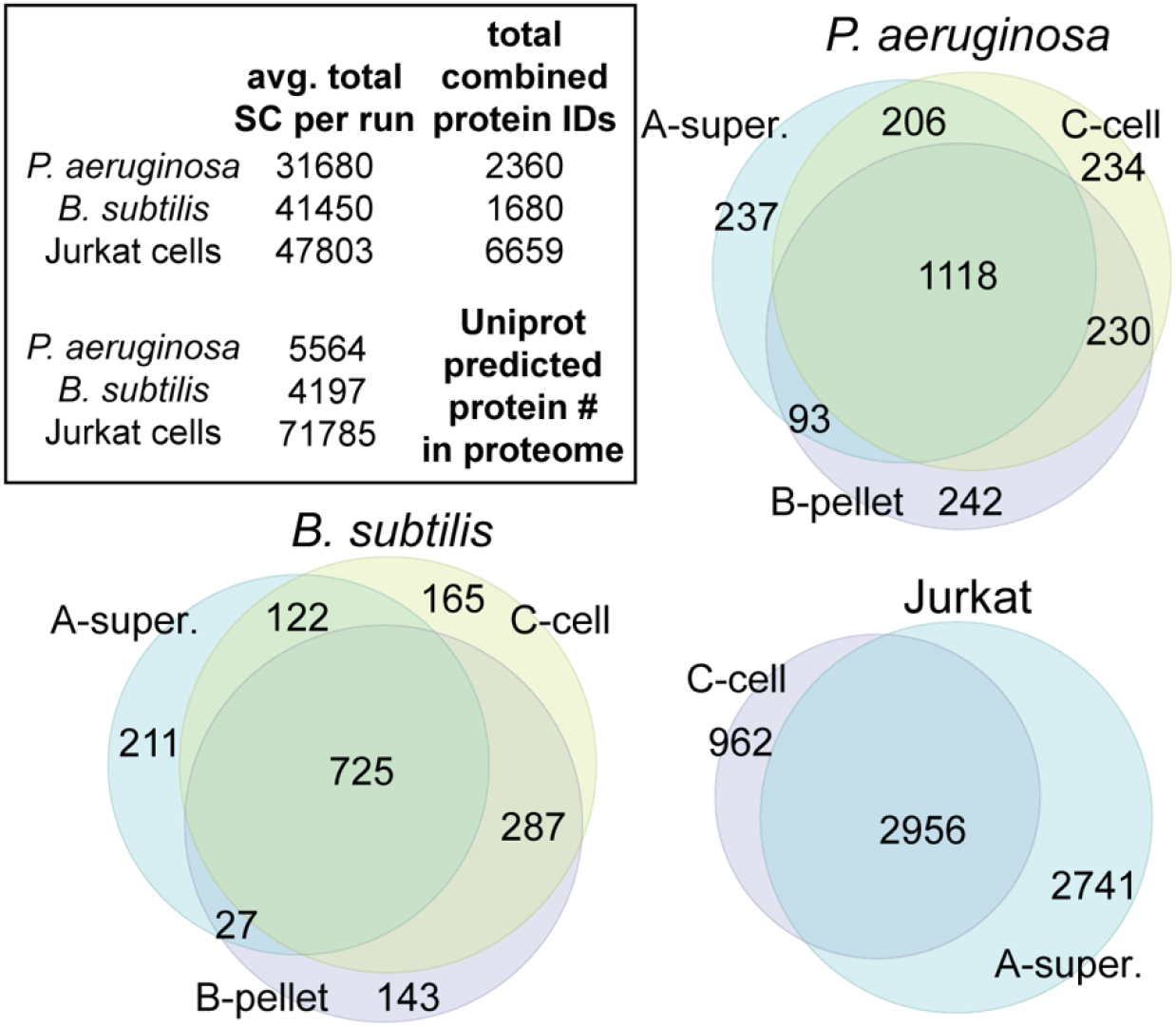
Venn diagrams depict number of proteins identified by LC-MS/MS for each treatment method pertaining to *P. aeruginosa, B. subtilis*, and human Jurkat cells. The total average peptide spectral matches (spectral counts, SC) per LC-MS/MS run (triplicate datasets for each treatment and three cell types), total combined proteins identified per cell type, and number of predicted proteins in respective organism’s genome are provided as a table in the inset.

**Figure 3.**
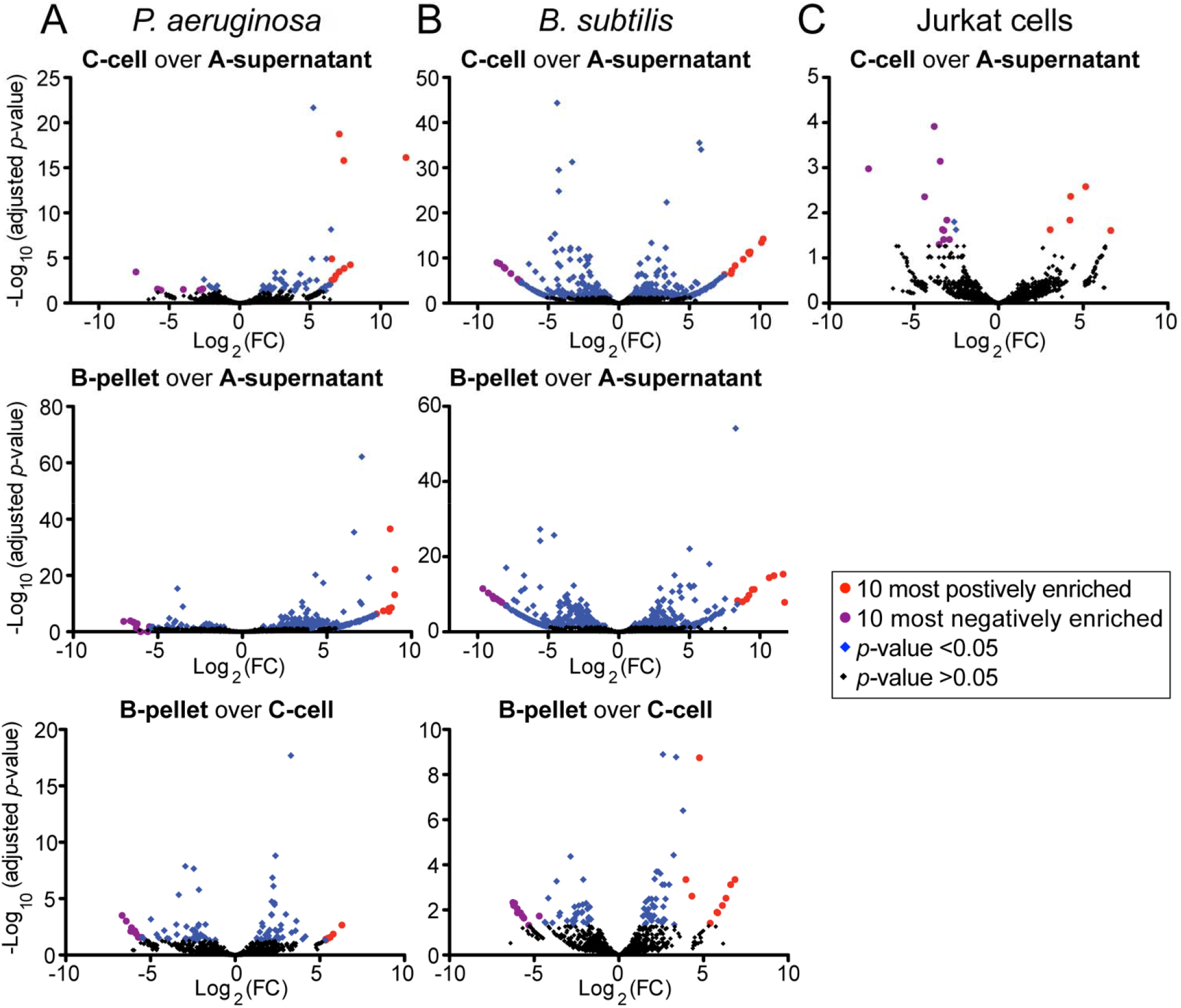
Volcano plots depict differentially detected proteins between treatment methods for *P. aeruginosa* (A), *B. subtilis* (B), and human Jurkat cells (C). Significantly enriched proteins (*p*-value <0.05) are colored blue. The top 10 positively and negatively detected proteins in the B-pellet are colored red and purple, respectively. A list of the 10 most significantly differentially detected proteins is available in the Supporting Information Tables 1-10 and the complete list of proteins is in the Supporting Information Dataset.

The detected quantities of 265 *P. aeruginosa* proteins shifted significantly between A-supernatant and B-pellet (Figure 3A). 209/264 proteins were positively enriched in B-pellet, with log_2_FC from 1.25 to 9.03 with TA treatment (Table S3). All of the 10 most-enriched proteins are annotated as envelope-associated, and 7 of these proteins were not detected in A-supernatant, including PhoP/Q and low Mg^2+^ inducible OM protein H1 (G3XD11), two probable OM proteins (Q9I456, Q9HVI2), motility protein FimV (Q9HZA6), lipid A deacylase PagL (Q9HVD1), membrane-bound lysozyme inhibitor (Q9I574), and OM protein assembly factor BamB (Q9HXJ7). 56/265 proteins were negatively enriched in B-pellet, ranging from log_2_FC –1.09 to –7.02 (Figure 3A). Of the 10 most shifted proteins, 4 are cytoplasmic, 4 are periplasmic, and two are of unknown localization (Table S4). For Gram^+^ *P. aeruginosa*, TA treatment favors the LC-MS/MS detection of envelope-associated proteins and clearly demonstrates that the post-sonication insoluble pellet contains important proteomic information missed with sonication-only preparations.

### Preparation of *B. subtilis* by sonication and TA treatment preparations yield complementary proteome information

The analysis of Gram^+^ *B. subtilis* identified a total of 1680 proteins, with 725 detected in all three sample preparations and 211, 165, and 143 unique proteins identified in A-supernatant, B-pellet, and C-cell, respectively (Figure 2). Between A-supernatant and C-cell datasets, 380 proteins were differentially detected, and 206/380 were found in greater abundance due to TA treatment, ranging from log_2_FC enrichments of 1.02 to 10.2 (Figure 3B). 6 of the 10 most-enriched proteins are cytoplasmic, including cyclo(L-leucyl-L-leucyl) synthase (O34351) and dimodular nonribosomal peptide synthase (P45745), and four are envelope-associated, including the pulcherriminic acid synthase (O34926) and tRNA nuclease WapA (Q07833) (Table S5). 174/380 proteins were negatively enriched in C-cell compared to A-supernatant. 3-ketoacyl-CoA thiolase (O32177) was the most differentially detected protein with a log_2_FC of –8.63 in C-cell(Table S6).

Differences between *B. subtilis* A-supernatant and B-pellet datasets were pronounced. 482 proteins were differentially detected, and 216/482 were positively enriched in B-pellet, with log_2_FC from 1.05 to 11.75 (Figure 3B). Similar to the comparison between A-supernatant and C-cell, a mixture of predicted cytoplasmic and envelope-associated proteins were differentially detected (Table S7). 266/482 differentially detected proteins were negatively enriched in B-pellet (log_2_FC −1.11 to -9.63), with glucose starvation-inducible protein B (P26907) as the most-changed (Table S8). In contrast to *P. aeruginosa*, most TA treatment-enriched *B. subtilis* proteins are not annotated as envelope-associated. However, the subcellular locales of *B. subtilis* proteins are not well annotated in general; thus, it is unclear in this case whether TA treatment broadly enables the enrichment of envelope-associated proteins. What is clear, however, is that sonication and TA treatment of *B. subtilis* resulted in datasets with complementary proteome information.

### Limited differences in Jurkat cell data between sonication and TA treatment preparations

A total of 6659 proteins were detected from Jurkat cells, with 2741 exclusive to A-supernatant, 962 in C-cell only, and 2956 common proteins (Figure 2). 17 proteins were differentially detected between the A-supernatant and C-cell preparations (Figure 3C). 5/17 were positively enriched, ranging from log_2_FC 3.05 to 6.63 in the C-cell dataset, and consisted of nucleic acid-binding proteins (Table S9). Conversely, 12/17 proteins were negatively shifted, ranging from log_2_FC −2.50 to −7.67 in C-cell, including cytoskeleton proteins and four isoforms of pyruvate kinase (Table S10). Overall, few Jurkat proteins with significant changes in detection were found; most proteins exclusive to A-supernatant versus C-cell were of low abundance. In addition, we suspect that TA treatment could be dissociating *N*-linked nucleobases from their sugar-phosphate backbones, thus disaggregating protein-nucleic acid complexes that are resistant to proteolytic cleavage under normal conditions. This, in turn, could explain the enrichment in detected nucleic acid binding proteins with TA treatment.

### TA treatment of human gut microbiome samples greatly improves protein detection

Metaproteomic analysis of a healthy human gut microbiome yielded a total of 8083 unique protein clusters^50^ collectively among the treatment methods, with an average of 22203 matched spectra per run. Principal component analysis (Figure 4A) and protein overlap demonstrate that B-pellet (7021 clusters) and C-cell (6796 clusters) datasets are most similar to each other, and both differ in detectable protein composition from A-supernatant (4569 clusters) (Figures 4B, 4C).^16^ Of the 8083 protein clusters identified, 5240 were differentially detected between A-supernatant and C-cell, with adjusted *p-*value <0.05 and |log_2_(FC)| >1 (Figure 5A). 3720/5240 differentially detected protein clusters were positively enriched in C-cell, and several of the most-enriched proteins included elongation factor Tu from a range of bacteria (Table S11). Conversely, 1520/5240 clusters diminished with respect to TA treatment (Table S12). 5180 protein clusters were differentially detected between A-supernatant and B-pellet with adjusted *p-*value <0.05 and |log_2_FC| >1 (Figure 5B). 3722/5180 were positively enriched with TA treatment, and 7 of the top 10 protein clusters were the same as those enriched in the C-cell versus A-supernatant (Table S13). 1458/5180 proteins were negatively enriched with TA treatment and vary in composition and parental bacterial species (Table S14). Interestingly, only 786 protein clusters were differentially detected between B-pellet (456/786) and C-cell (330/786) (Figure 5C), and two of the most positively enriched clusters in B-pellet were human (Tables S15 and S16). Surprisingly, additional human proteins were identified in the B-pellet dataset despite extensive washing prior to bacterial lysis. Though TA treatment enables the detection of many new microbiome proteins, our subcellular localization analysis is limited by the poorly characterized nature of the gut microbiome’s proteome (Figures 5D-5G).^55,56^

**Figure 4.**
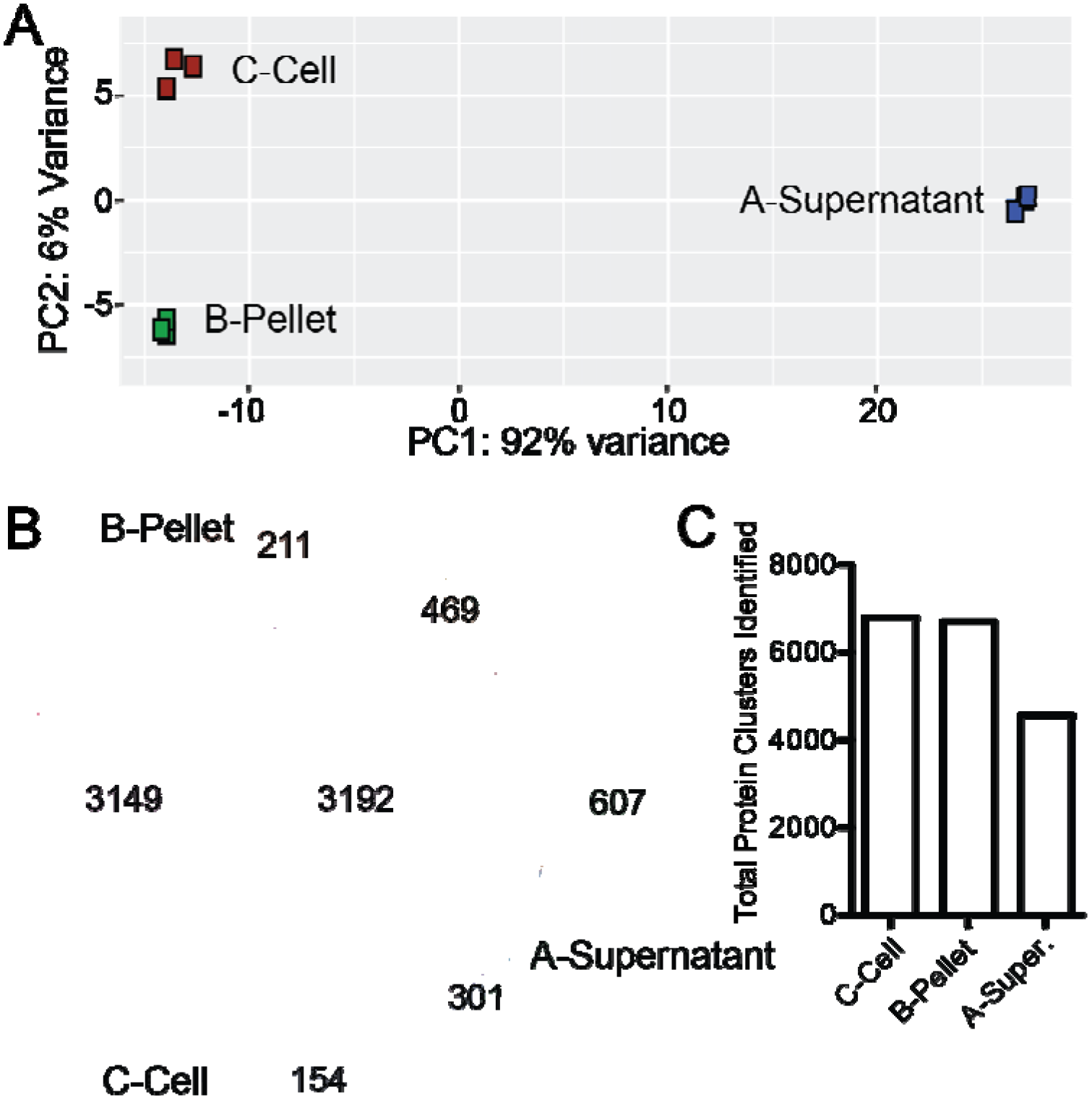
Analysis of single human microbiome sample by three treatment methods. (A) Principle component analysis of LC-MS/MS-detected microbiome proteome by three treatment methods. (B) Venn diagram depicti numbers of protein clusters identified by LC-MS/MS for each treatment method. (C) Total number of protein clusters identified per treatment group.

**Figure 5.**
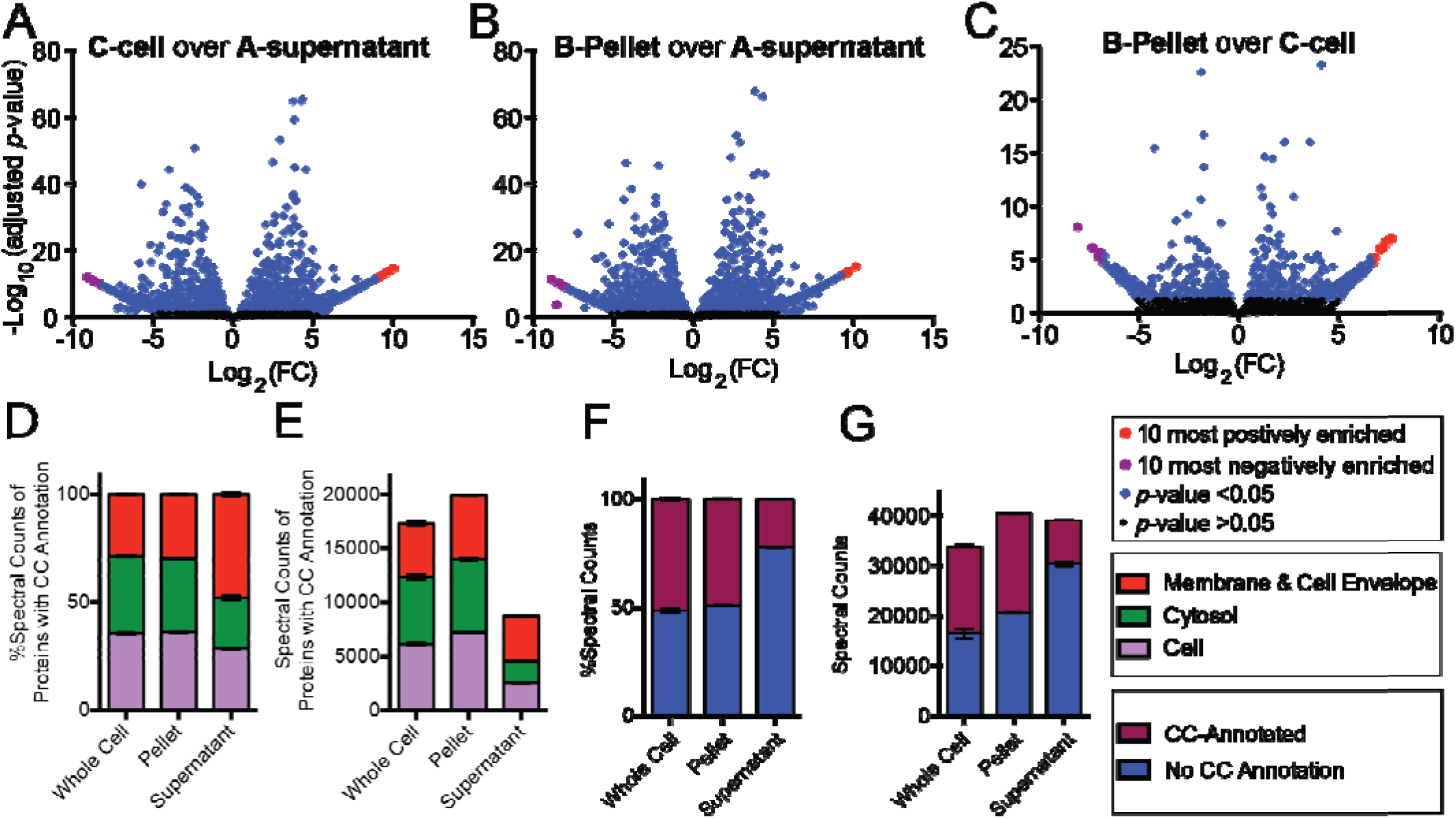
Volcano plots of altered protein clusters in C-cell compared to A-supernatant (A), B-pellet compared A-supernatant (B), and B-pellet compared to C-cell (C) for human microbiome samples. Protein clusters significantly enriched (*p*-value <0.05) are colored blue, and the top 10 positively and negatively enriched prot clusters for each volcano plot are colored red and purple, respectively. A list of the 10 most significantly differentially detected proteins is available in the Tables S11-S16 and the complete list of proteins is in the Supporting Information Dataset. (D, E) Cellular component GO term classification of LC-MS/MS detected microbiome proteins by treatment method. (F, G) Comparison of detected microbiome proteins with and wit cellular component GO term annotations.

One of the most remarkable outcomes of our investigation was the impact of TA treatment on the microbiome A-supernatant fraction. LC-MS/MS analysis of the A-supernatant microbiome prepared similarly to the other bacterial and Jurkat cell A-supernatant samples returned 0 protein cluster matches despite .RAW files of adequate size and chromatographic quality (Figure S2). This is in sharp contrast to the B-pellet and C-cell microbiome dataset analyses that returned thousands of protein cluster matches. We hypothesized that the abundant unmatched features in microbiome A-supernatant could be non-peptide contaminants that are minimized in B-pellet and C-cell preparations by TA treatment. To test this hypothesis, we TA-treated the same microbiome A-supernatant sample and gratifyingly observed >2000 protein cluster matches after LC-MS/MS analysis, a conspicuous difference compared to the 0 protein cluster matches we observed before TA treatment.

Problems related to ion suppression and signal interference by non-peptide contaminants in protein preparations have been documented and addressed in previous studies.^17,19–21,23,25,28,32,36,38^ For example, with regard to metaproteomic surveys of the soil microbiota, humic acid-type substances have been identified as prominent interfering contaminants. Humic acids have diverse chemical structures and are particularly difficult to completely remove.^38^ For human microbiome samples, it is unclear at this time whether interfering compounds fit a particular chemical profile but it appears that TA treatment markedly ameliorates their impact on microbiome analysis.

## CONCLUSIONS

Use of TA in the preparation of bacteria dramatically impacts the detectable proteome. Many proteins were uniquely detected by both sonication and TA treatment, clearly demonstrating the complementarity of TA treatment to established preparation methods. For the Gram^-^ *P. aeruginosa*, TA treatment enables the enrichment of envelope-associated proteins by LC-MS/MS. For the Gram^+^ *B. subtilis*, a profound enrichment of many proteins is observed, including many envelope-associated proteins; however, there are also many proteins that only appear to be detected with sonication, demonstrating the complementarity of TA treatment to other preparation methods. Finally, TA treatment not only appears robust for a microbiome sample, but it also enabled the analysis of supernatant prepared by sonication that otherwise yielded a null dataset. We anticipate that TA treatment will have broad applicability for the analysis of complex microbial systems.

## ASSOCIATED CONTENT

“Wang et al 2018 SI Dataset.xlsx”contains the full list of detected proteins from each LC-MS/MS data set.

Supplementary tables and figures can be found in “Wang et al 2018 SI.pdf”

**Figure S1.** Detailed schematic overview of two proteome preparation methods.

**Figure S2.** Base-peak chromatograms for microbiome A-supernatant samples.

**Table S1.** 10 most positively enriched proteins in *P. aeruginosa* C-cell dataset relative to A-supernatant.

**Table S2.** 10 most negatively enriched proteins in *P. aeruginosa* C-cell dataset relative to A-supernatant.

**Table S3.** 10 most positively enriched proteins in *P. aeruginosa* B-pellet dataset relative to A-supernatant.

**Table S4.** 10 most negatively enriched proteins in *P. aeruginosa* B-pellet dataset relative to A-supernatant.

**Table S5.** 10 most positively enriched proteins in *B. subtilis* C-cell dataset relative to A-supernatant.

**Table S6.** 10 most negatively enriched proteins in *B. subtilis* C-cell dataset relative to A-supernatant.

**Table S7.** 10 most positively enriched proteins in *B. subtilis* B-pellet dataset relative to A-supernatant.

**Table S8.** 10 most negatively enriched proteins in *B. subtilis* B-pellet dataset relative to A-supernatant.

**Table SI 9** Positively enriched proteins in human Jurkat cells C-cell dataset relative to A-supernatant.

**Table S10.** Negatively enriched proteins in human Jurkat cells C-cell dataset relative to A-supernatant.

**Table S11.** 10 most positively enriched protein clusters in human microbiome C-cell dataset relative to Asupernatant.

**Table S12.** 10 most negatively enriched protein clusters in human microbiome C-cell dataset relative to Asupernatant.

**Table S13.** 10 most positively enriched protein clusters in human microbiome B-pellet dataset relative to Asupernatant.

**Table S14.** 10 most negatively enriched protein clusters in human microbiome B-pellet dataset relative to Asupernatant.

**Table S15.** 10 most positively enriched protein clusters in human microbiome B-pellet dataset relative to C-cell.

**Table S16.** 10 most negatively enriched protein clusters in human microbiome B-pellet dataset relative to C-cell.

## AUTHOR INFORMATION

### Author Contributions

A.Y.W., P.S.T.-B., and D.W.W. conceived of the project. P.S.T.-B. and A.Y.W. performed wet-lab experiments and collected MudPIT LC-MS/MS data. A.Y.W., P.S.T.-B., and G.S.S. analyzed tandem LC-MS/MS data. A.Y.W., P.S.T.-B., and G.S.S. contributed to peptide mapping and functional analysis. The manuscript was written through contributions of all authors. All authors have given approval to the final version of the manuscript.

### Funding Sources

We also gratefully acknowledge financial support from The Scripps Research Institute, Boehringer Ingelheim (to D.W.W. and A.I.S.), U54GM114833 (to A.I.S.), and US Environmental Protection Agency STAR Pre-doctoral Fellowship FP917296-01-0 (to P.S.T.-B.)

### Notes

The authors declare that they have no competing interests.

## ACKNOWLEDGEMENTS

We thank Dr. J. Yates, Dr. J. Moresco, and Dr. J. Diedrich for technical assistance with mass spectrometry instrumentation; Dr. M. Perego for providing the *B. subtilis* strain.

